# Inhibitory control of correlated intrinsic variability in cortical networks

**DOI:** 10.1101/041103

**Authors:** Carsen Stringer, Marius Pachitariu, Michael Okun, Peter Bartho, Kenneth Harris, Peter Latham, Maneesh Sahani, Nicholas Lesica

## Abstract

Cortical networks exhibit intrinsic dynamics that drive coordinated, large-scale fluctuations across neuronal populations and create noise correlations that impact sensory coding. To investigate the network-level mechanisms that underlie these dynamics, we developed novel computational techniques to fit a deterministic spiking network model directly to multi-neuron recordings from different species, sensory modalities, and behavioral states. The model generated correlated variability without external noise and accurately reproduced the wide variety of activity patterns in our recordings. Analysis of the model parameters suggested that differences in noise correlations across recordings were due primarily to differences in the strength of feedback inhibition. Further analysis of our recordings confirmed that putative inhibitory neurons were indeed more active during desynchronized cortical states with weak noise correlations. Our results demonstrate that network models with intrinsically-generated variability can accurately reproduce the activity patterns observed in multi-neuron recordings and suggest that inhibition modulates the interactions between intrinsic dynamics and sensory inputs to control the strength of noise correlations.

## Introduction

The patterns of cortical activity evoked by sensory stimuli provide the internal representation of the outside world that underlies perception. However, these patterns are driven not only by sensory inputs, but also by the intrinsic dynamics of the underlying cortical network. These dynamics can create correlations in the activity of neuronal populations with important consequences for coding and computation^1–3^. The correlations between pairs of neurons have been studied extensively^3–5^, and recent studies have demonstrated that they are driven by dynamics involving coordinated, large-scale fluctuations in the activity of many cortical neurons^6–8^. Inactivation of the cortical circuit nearly completely quenches these synchronized fluctuations in both awake and anesthetized animals, suggesting that this synchronization is cortical in origin^9^. Importantly, the nature of these dynamics and the correlations that they create are dependent on the state of the underlying network; it has been shown that various factors modulate the strength of correlations, such as anaesthesia^10–12^, attention^13–15^, locomotion^16,17^, and alertness^18,19^. In light of these findings, it is critical that we develop a deeper understanding of the origin and coding consequences of correlations at the biophysical network level.

While a number of modeling studies have explored the impact of correlations on sensory coding^1,3,20–23^, there have been few efforts to identify their biophysical origin; the standard assumption that correlations arise from common input noise^20,24,25^ simply pushes the correlations from spiking to the membrane voltage without providing insight into their genesis. Models that use external noise to create correlations have been used in theoretical investigations of how network dynamics can transform correlations^24^, but no physiological source for the external noise used in these models has yet been identified. However, no external noise is needed to generate the correlated activity that is observed in vivo; in vitro experimental studies have shown that cortical networks are capable of generating large-scale fluctuations intrinsically^26,27^, and in vivo results suggest that the majority of cortical fluctuations arise locally^9^. If the major source of the correlations in cortical networks is, in fact, internal, then the network features that control these correlations may be different from those that control correlations in model networks with external noise.

We demonstrate here that network models with intrinsic variability are indeed capable of reproducing the activity patterns that are observed in vivo, and then proceed to use a large number of multi-neuron recordings and a model-based analysis to investigate the mechanisms that control intrinsically generated-noise correlations. For our results to provide direct insights into physiological mechanisms, we required a model with several properties: (1) the model must be able to internally generate the complex intrinsic dynamics of cortical networks, (2) it must be possible to fit the model parameters directly to spiking activity from individual multi-neuron recordings, and (3) the model must be biophysically interpretable and enable predictions that can be tested experimentally. No existing model satisfies all of these criteria; the only network models that have been fit directly to multi-neuron recordings have relied on either abstract dynamical systems^28^ or probabilistic frameworks in which variability is modelled as stochastic and correlated variability arises through abstract latent variables whose origin is assumed to lie either in unspecified circuit processes^21,29–31^ or elsewhere in the brain^20,32^. While these models are able to accurately reproduce many features of cortical activity and provide valuable summaries of the phenomenological and computational properties of cortical networks, their parameters are difficult to interpret at a biophysical level.

One alternative to these abstract stochastic models is a biophysical spiking network,^33–37^. These networks can be designed to have interpretable parameters, but have not been shown to internally generate large-scale fluctuations and noise correlations of the kind routinely seen in multi-neuron recordings. Networks with structured connectivity have been shown to generate correlated activity in small groups containing less than 5% of all neurons^36^, but not in the entire network. Furthermore, large-scale neural network models have not yet been fit directly to multi-neuron recordings and, thus, their use has been limited to attempts to explain qualitative features of cortical dynamics through manual tuning of network parameters. This inability to fit the networks directly to recordings has made it difficult to identify which of these network features, if any, play an important role in vivo. To overcome this limitation, we used a novel computational approach that allowed us to fit spiking networks directly to individual multi-neuron recordings. By taking advantage of the computational power of graphics processing units (GPUs), we were able to simulate the network with millions of different parameter values for 800 seconds each to find those that best reproduced the structure of the activity in a given recording.

We developed a novel biophysical spiking network with intrinsic variability and a small number of parameters that was able to capture the apparently “doubly chaotic” structure of cortical activity^38^. Like classical excitatory-inhibitory networks, the model generates deterministic “microscopic” trial-to-trial variability in the spike times of individual neurons^33^, as well as “macroscopic” variability in the form of coordinated, large-scale fluctuations that are shared across neurons. Because these fluctuations are of variable duration, arise at random times, and do not necessarily phase-lock to external input, they create noise correlations in evoked responses. This intrinsic variability distinguishes our model from previous rate or spiking network models^24,35,37^, as well as from phenomenological dynamical systems^30,31^, all of which create noise correlations by injecting common noise into all neurons (an approach which, by construction, provides little insight into the biophysical mechanisms that generate the noise^24^).

To gain insight into the mechanisms that control noise correlations in vivo, we took the following approach: (1) we assembled multi-neuron recordings from different species, sensory modalities, and behavioral states to obtain a representative sample of cortical dynamics; (2) we generated activity from the network model to understand how each of its parameters controls its dynamics, and we verified that it was able to produce a variety of spike patterns that were qualitatively similar to those observed in vivo; (3) we fit the model network directly to the spontaneous activity in each of our recordings, and we verified that the spike patterns generated by the network quantitatively matched those in each recording; (4) we examined responses to sensory stimuli to determine which of the model parameters could account for the differences in noise correlations across recordings – the results of this analysis identified the strength of feedback inhibition as a key parameter and predicted that the activity of inhibitory interneurons should vary inversely with the strength of noise correlations; (5) we confirmed this prediction through additional analysis of our recordings showing that the activity of putative inhibitory neurons is increased during periods of cortical desynchronization with weak noise correlations in both awake and anesthetized animals. Our results suggest that weak inhibition allows activity to be dominated by coordinated, large-scale fluctuations that cause the state of the network to vary over time and, thus, create variability in the responses to successive stimuli that is correlated across neurons. In contrast, when inhibition is strong, these fluctuations are suppressed and the network state remains constant over time, allowing the network to respond reliably to successive stimuli and eliminating noise correlations.

## Results

### Cortical networks exhibit a wide variety of intrinsic dynamics

To obtain a representative sample of cortical activity patterns, we collected multi-neuron recordings from different species (mouse, gerbil, or rat), sensory modalities (A1 or V1), and behavioral states (awake or under one of several anesthetic agents). We compiled recordings from a total of 59 multi-neuron populations across 6 unique recording types (i.e. species/modality/state combinations; see Supplementary Table 1). The spontaneous activity in different recordings exhibited striking differences not only in overall activity level, but also in the spatial and temporal structure of activity patterns; while concerted, large-scale fluctuations were prominent in some recordings, they were nearly absent in others (Figure 1a). In general, large-scale fluctuations were weak in awake animals and strong under anesthesia, but this was not always the case (see further examples in Figure 3 and summary statistics for each recording in Supplementary Figure 1).

**Figure 1.**
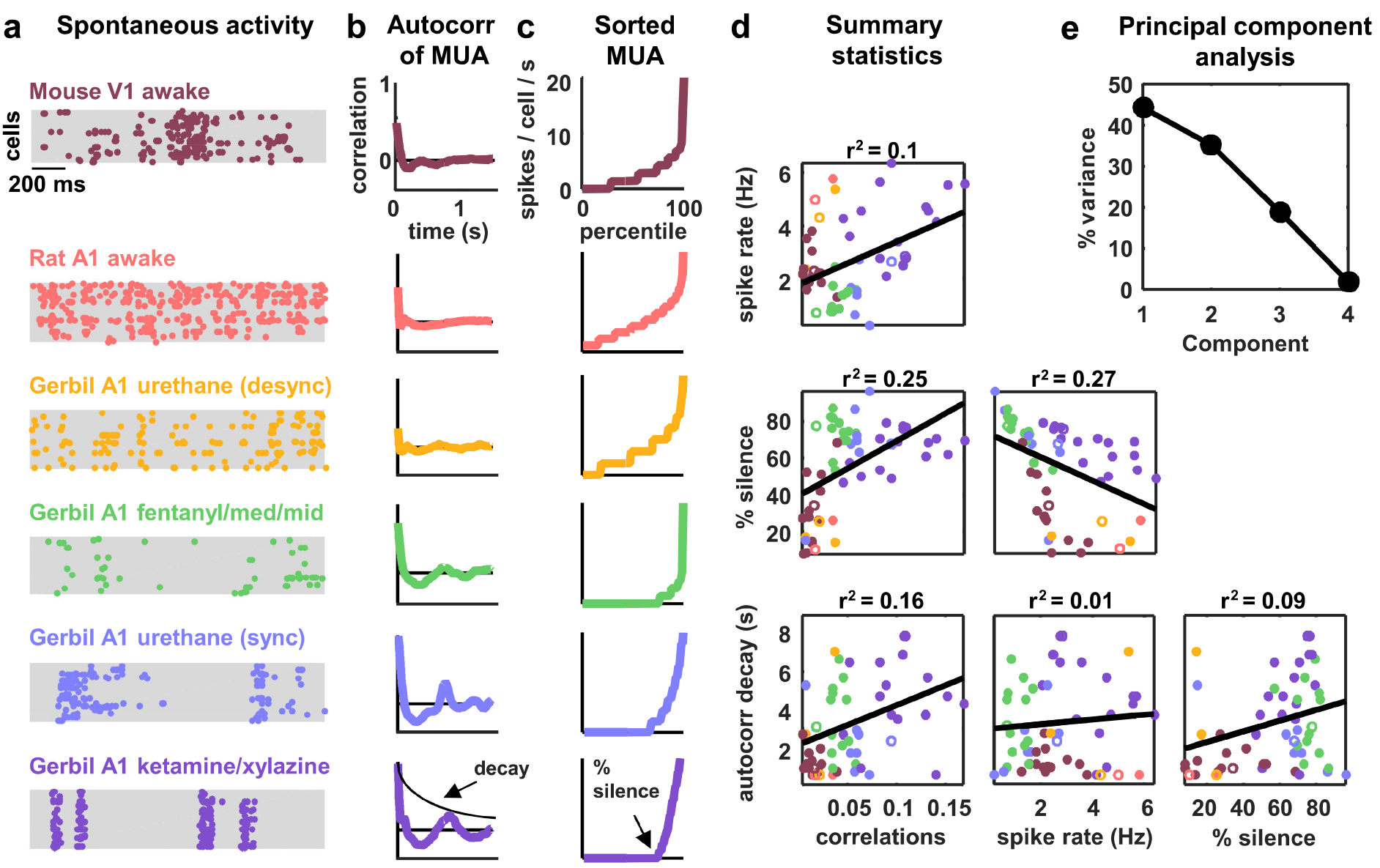
**Cortical networks exhibit a wide variety of intrinsic dynamics** (a) Multi-neuron raster plots showing examples of a short segment of spontaneous activity from each of our recording types. Each row in each plot represents the spiking of one single unit. Note that recordings made under urethane were separated into two different recording types, synchronized (’sync’) and desynchronized (’desync’), as described in the Methods. (b) The autocorrelation function of the multi-unit activity (MUA, the summed spiking of all neurons in the population in 15 ms time bins) for each example recording. The timescale of the autocorrelation function (the ‘autocorr decay’) was measured by fitting an exponential function to its envelope as indicated. (c) The values of the MUA across time bins sorted in ascending order. The percentage of time bins with zero spikes (the ‘% silence’) is indicated. (d) Scatter plots showing all possible pairwise combinations of the summary statistics for each recording. Each point represents the values for one recording. Colors correspond to recording types as in A. The recordings shown in A are denoted by open circles. The best fit line and the fraction of the variance that it explained are indicated on each plot. Spearman’s rank correlation p-values for each plot (from left to right, top to bottom) are as follows: p<0.05, p<10^−4^, p<10^−5^, p<10^−2^, p=0.447, p<0.05. (e)The percent of the variance in the summary statistics across recordings that is explained by each principal component of the values.

The magnitude and frequency of the large-scale fluctuations in each recording were reflected in the autocorrelation function of the multi-unit activity (MUA, the summed spiking of all neurons in the population in 15 ms time bins). The autocorrelation function of the MUA decayed quickly to zero for recordings with weak large-scale fluctuations, but had oscillations that decayed slowly for recordings with stronger fluctuations (Figure 1b). The activity patterns in recordings with strong large-scale fluctuations were characterized by clear transitions between up states, where most of the population was active, and down states, where the entire population was silent. These up and down state dynamics were reflected in the distribution of the MUA across time bins; recordings with strong large-scale fluctuations had a large percentage of time bins with zero spikes (Figure 1c).

To summarize the statistical structure of the activity patterns in each recording, we measured four quantities. We used mean spike rate to describe the overall level of activity, mean pairwise correlations to describe the spatial structure of the activity patterns, and two different measures to describe the temporal structure of the activity patterns – the decay time of the autocorrelation function of the MUA, and the percentage of MUA time bins with zero spikes. While there were some dependencies in the values of these quantities across different recordings (Figure 1d), there was also considerable scatter both within and across recording types. This scatter suggests that there is no single dimension in the space of cortical dynamics along which the overall level of activity and the spatial and temporal structure of the activity patterns all covary, but rather that cortical dynamics span a multi-dimensional continuum^10^. This was confirmed by principal component analysis; even in the already reduced space described by our summary statistics, three principal components were required to account for the differences in spike patterns across recordings (Figure 1e).

### A deterministic spiking network model of cortical activity

To investigate the network-level mechanisms that control cortical dynamics, we developed a biophysically-interpretable model that was capable of reproducing the wide range of activity patterns observed in vivo. We constructed a minimal deterministic network of excitatory spiking integrate-and-fire neurons with non-selective feedback inhibition and single-neuron adaptation currents (Figure 2a). Each neuron receives constant tonic input, and the neurons are connected randomly and sparsely with 5% probability. The neurons are also coupled indirectly through global, supralinear inhibitory feedback driven by the spiking of the entire network^39^, reflecting the near-complete interconnectivity between pyramidal neurons and interneurons in local populations^40–42^. The supralinearity of the inhibitory feedback is a critical feature of the network, as it shifts the balance of excitation and inhibition in favor of inhibition when the network is strongly driven, as has been observed in awake animals^43^.

The model has five free parameters: three controlling the average strength of excitatory connectivity, the strength of inhibitory feedback, and the strength of adaptation, respectively, and two controlling the strength of the tonic input to each neuron, which is chosen from an exponential distribution. The timescales that control the decay of the excitatory, inhibitory and adaptation currents are fixed at 5 ms, 3.75 ms and 375 ms, respectively. (These timescales have been chosen based on the physiologically known timescales of AMPA, GABA_a_, and the calcium-dependent afterhyperpolarizing current. We also verified that the qualitative nature of our results did not change when we included slow conductances or clustered connectivity; see Supplementary Figure 2).

Note that no external noise input is required to generate variable activity; population-wide fluctuations over hundreds of milliseconds are generated when the slow adaptation currents synchronize across neurons to maintain a similar state of adaptation throughout the entire network, which, in turn, results in coordinated spiking^44,45^. The variability in the model arises through chaotic amplification of small changes in initial conditions or small perturbations to the network that cause independent simulations to diverge. In some parameter regimes, the instability of the network is such that the structure of the spike patterns generated by the model is sensitive to changes in the spike times of individual neurons. In fact, a single spike added randomly to a single neuron during simulated activity is capable of changing the time course of large-scale fluctuations, in some cases triggering immediate population-wide spiking (Figure 2b, top rows). Similar phenomena have been observed in vivo previously^46^ and were also evident in our recordings when comparing different extracts of cortical activity; spike patterns that were similar for several seconds often then began to diverge almost immediately (Figure 2b, bottom rows).

### Multiple features of the network model can control its dynamics

The dynamical regime of the network model is determined by the interactions between its different features. To determine the degree to which each feature of the network was capable of influencing the structure of its activity patterns, we analyzed the effects of varying the value of each model parameter. We started from a fixed set of parameter values and simulated activity while independently sweeping each parameter across a wide range of values. The results of these parameter sweeps clearly demonstrate that each of the five parameters can exert strong control over the dynamics of the network, as both the overall level of activity and the spatial and temporal structure of the patterns in simulated activity varied widely with changes in each parameter (Figure 2c–d).

With the set of fixed parameter values used for the parameter sweeps, the network is in a regime with slow, ongoing fluctuations between up and down states. In this regime, the amplification of a small perturbation results in a sustained, prolonged burst of activity (up state), which, in turn, drives a build-up of adaptation currents that ultimately silences the network for hundreds of milliseconds (down state) until the cycle repeats. These fluctuations can be suppressed by an increase in the strength of feedback inhibition, which eliminates slow fluctuations and shifts the network into a regime with weak, tonic spiking and weak correlations (Figure 2c–d, first column); in this regime, small perturbations are immediately offset by the strong inhibition and activity is returned to baseline. Strong inhibition also offsets externally-induced perturbations in balanced networks^35^, but in our model such perturbations are internally-generated and would result in runaway excitation in the absence of inhibitory stabilization. The fluctuations between up and down states can also be suppressed by decreasing adaptation (Figure 2c–d, second column); without adaptation currents to create slow, synchronous fluctuations across the network, neurons exhibit strong, tonic spiking.

The dynamics of the network can also be influenced by changes in the strength of the recurrent excitation or tonic input. Increasing the strength of excitation results in increased activity and stronger fluctuations, as inhibition is unable to compensate for the increased amplification of small perturbations (Figure 2c–d, third column). Increasing the spread or baseline level of tonic input also results in increased activity, but with suppression, rather than enhancement, of slow fluctuations (Figure 2c–Figure 2d, fourth and fifth column). As either the spread or baseline level of tonic input is increased, more neurons begin to receive tonic input that is sufficient to overcome their adaptation current and, thus, begin to quickly reinitiate up states after only brief down states and, eventually, transition to tonic spiking.

**Figure 2.**
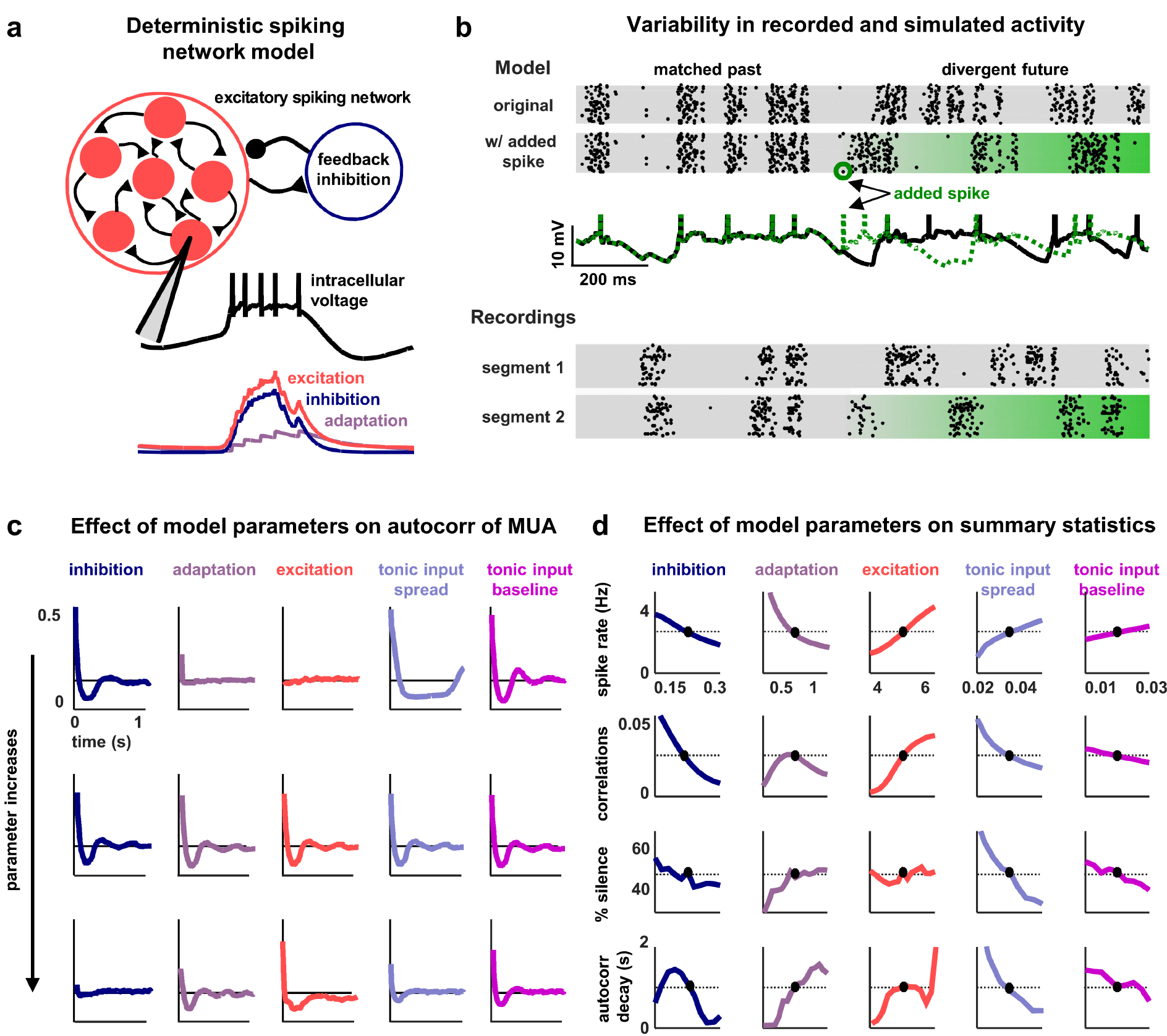
**A deterministic spiking network model of cortical activity** (a) A schematic diagram of our deterministic spiking network model. An example of a short segment of the intracellular voltage of a model neuron is also shown, along with the corresponding excitatory, inhibitory and adaptation currents. (b) An example of macroscopic variability in cortical recordings and network simulations. The top two multi-neuron raster plots show spontaneous activity generated by the model. By adding a very small perturbation, in this case one spike added to a single neuron, the subsequent activity patterns of the network can change dramatically. The middle traces show the intracellular voltage of the model neuron to which the spike was added. The bottom two raster plots show a similar phenomenon observed in vivo. Two segments of activity extracted from different periods during the same recording were similar for three seconds, but then immediately diverged. (c) The autocorrelation function of the MUA measured from network simulations with different model parameter values. Each column shows the changes in the autocorrelation function as the value of one model parameter is changed while all others are held fixed. The fixed values used were w_I_ = 0.22, w_A_ = 0.80, w_E_ = 4.50, b_1_ = 0.03, b_0_ = 0.013. (d) The summary statistics measured from network simulations with different model parameter values. Each line shows the changes in the indicated summary statistic as one model parameter is changed while all others are held fixed. Fixed values were as in panel c.

### The network model reproduces the dynamics observed in vivo

The network simulations demonstrate that each of its features is capable of controlling its dynamics and shaping the structure of its activity patterns. To gain insight into the mechanisms that may be responsible for creating the differences in dynamics observed in vivo, we fit the model to each of our recordings. We optimized the model parameters so that the patterns of activity generated by the network matched those observed in spontaneous activity (Figure 3a). We measured the agreement between the simulated and recorded activity by a cost function which was the sum of discrepancies in the autocorrelation function of the MUA, the distribution of MUA values across time bins, and the mean pairwise correlations. Together, these statistics describe the overall level of activity in each recording, as well as the spatial and temporal structure of its activity patterns.

Fitting the model to the recordings required us to develop new computational techniques. The network parametrization is fundamentally nonlinear, and the statistics used in the cost function are themselves nonlinear functions of a dynamical system with discontinuous integrate-and-fire mechanisms. Thus, as no gradient information was available to guide the optimization, we used Monte Carlo simulations to generate activity and measure the relevant statistics with different parameter values. By using GPU computing resources, we were able to design and implement network simulations that ran 10000× faster than real time, making it feasible to sample the cost function with high resolution and locate its global minimum to identify the parameter configuration that resulted in activity patterns that best matched those of each recording. We also verified that the global minimum of the cost function could be identified with 10× fewer samples of simulated activity using a Gibbs sampling optimizer with simulated annealing (Supplementary Figure 3), but the results presented below are based on the global minima identified by the complete sampling of parameter space.

The model was flexible enough to capture the wide variety of activity patterns observed across our recordings, producing both decorrelated, tonic spiking and coordinated, large-scale fluctuations between up and down states as needed (see examples in Figure 3b, statistics for all recordings and models in Supplementary Figure 1, and parameter values and goodness-of-fit measures for all recordings in Supplementary Figure 4). Because we used a cost function that captured many different properties of the recorded activity while fitting only a very small number of model parameters, the risk of network degeneracies was relatively low^47,48^. Nonetheless, we also confirmed that analysis of model parameters corresponding to local minima of the cost function did not lead to a different interpretation of our results (see Supplementary Figure 5).

**Figure 3.**
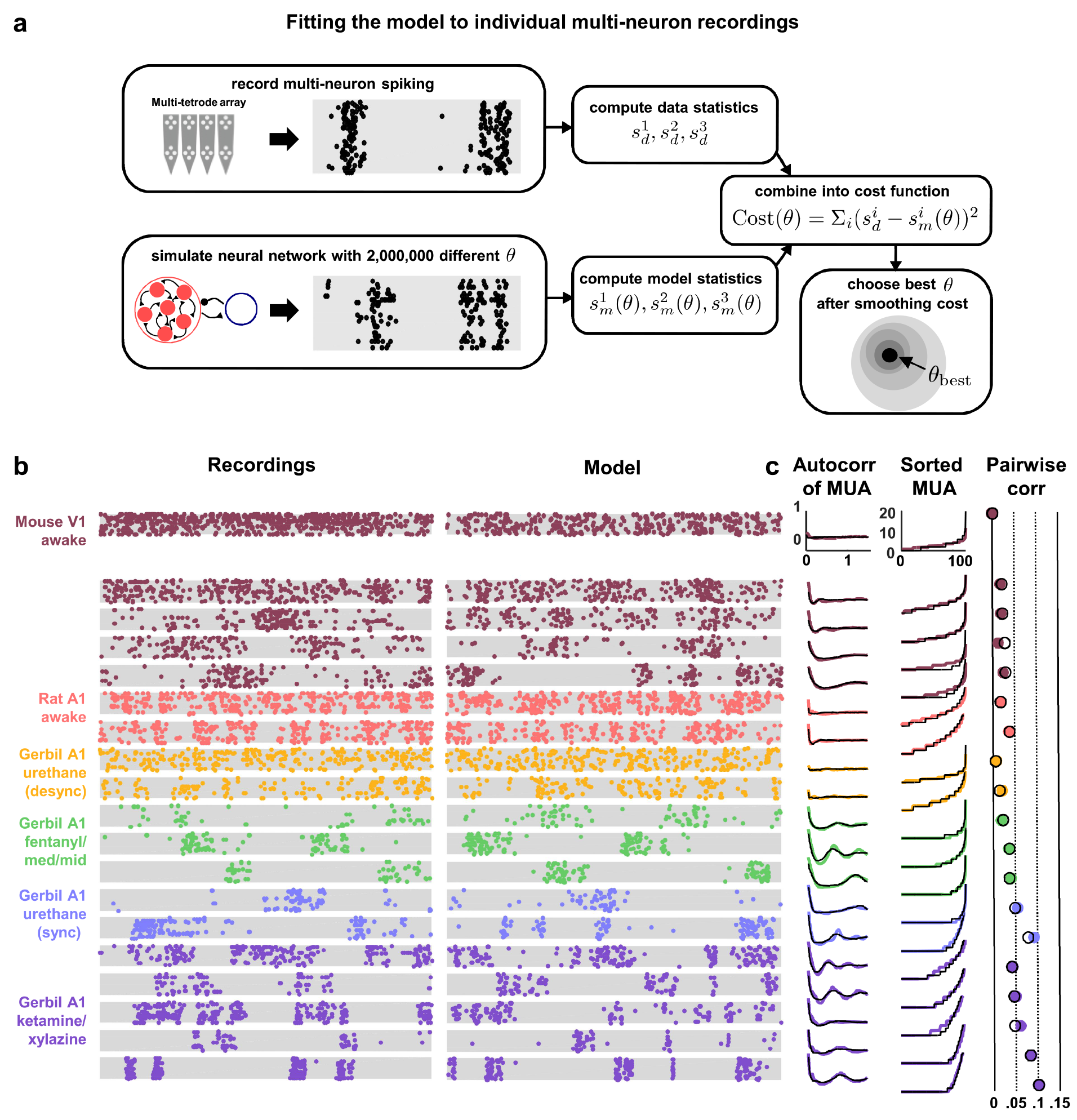
**Deterministic spiking networks reproduce the dynamics observed in vivo** (a) A schematic diagram illustrating how the parameters of the network model were fit to individual multi-neuron recordings. (b) Examples of spontaneous activity from different recordings, along with spontaneous activity generated by the model fit to each recording. (c) The left column shows the autocorrelation function of the MUA for each recording, plotted as in Figure 1. The black lines show the autocorrelation function measured from spontaneous activity generated by the model fit to each recording. The middle column shows the sorted MUA for each recording along with the corresponding model fit. The right column shows the mean pairwise correlations between the spiking activity of all pairs of neurons in each recording (after binning activity in 15 ms bins). The colored circles show the correlations measured from the recordings and the black open circles show the correlations measured from from spontaneous activity generated by the model fit to each recording.

**Figure 4.**
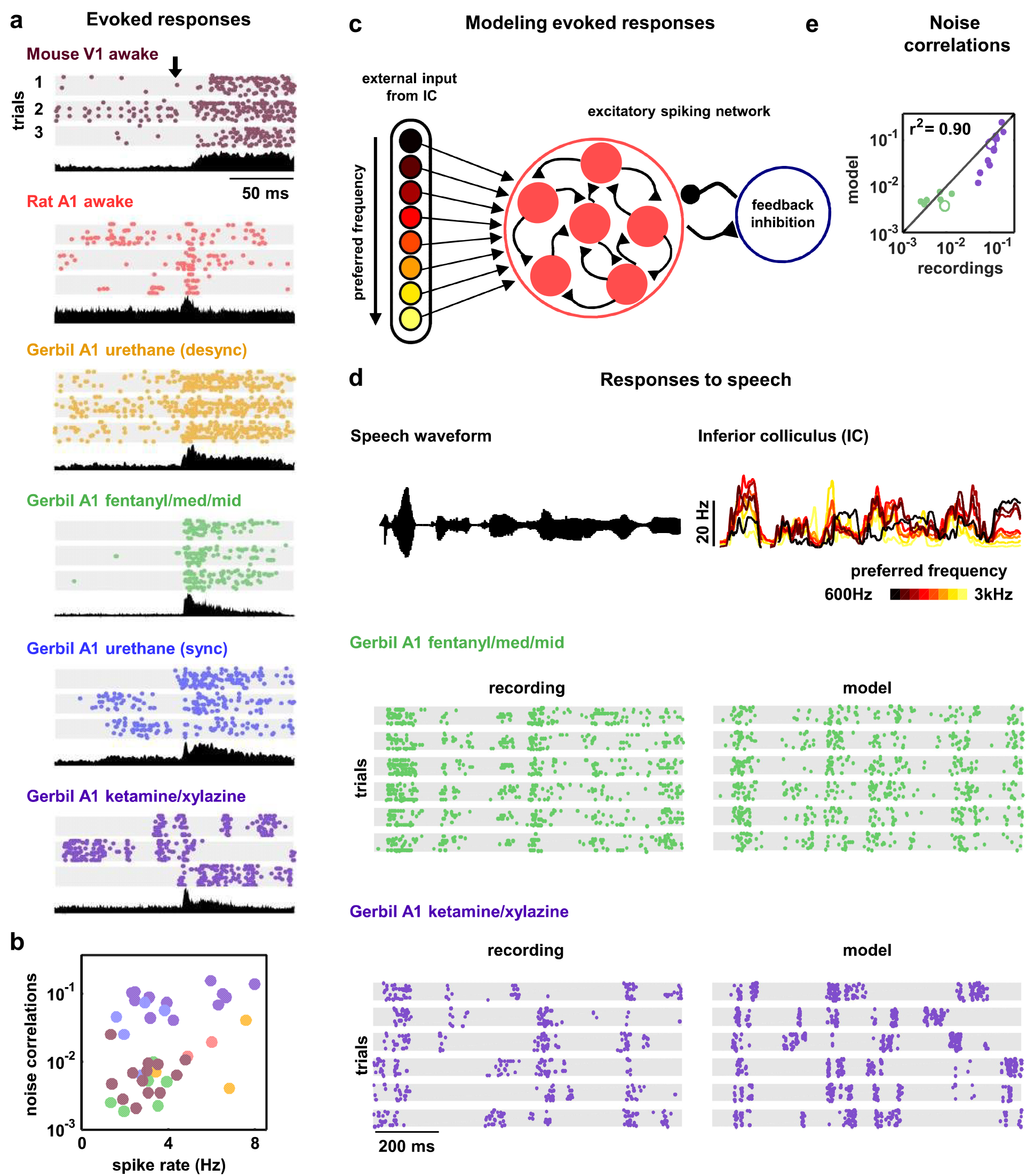
**Deterministic spiking networks reproduce the noise correlations observed in vivo** (a) Multi-neuron raster plots and PSTHs showing examples of evoked responses from each of our recording types. Each row in each raster plot represents the spiking of one single unit. Each raster plot for each recording type shows the response on a single trial. The PSTH shows the MUA averaged across all presentations of the stimulus. Different stimuli were used for different recording types (see Methods). (b) A scatter plot showing the mean spike rates and mean pairwise noise correlations (after binning the evoked responses in 15 ms bins) for each recording. Each point represents the values for one recording. Colors correspond to recording types as in panel a. The recordings shown in panel a are denoted by open circles. Values are only shown for the 38 of 59 recordings that contained both spontaneous activity and evoked responses. The Spearman’s rank correlation was significant with p=0.0105. (c) A schematic diagram illustrating the modelling of evoked responses. We constructed the external input using recordings of responses from more than 500 neurons in the inferior colliculus (1C), the primary relay nucleus of the auditory midbrain that provides the main input to the thalamocortical circuit. We have shown previously that the Fano factors of the responses of 1C neurons are close to 1 and the noise correlations between neurons are extremely weak (Garcia-Lazaro et al, 2013), suggesting that the spiking activity of a population of 1C neurons can be well described by series of independent, inhomogeneous Poisson processes. To generate the responses of each model network to the external input, we averaged the activity of each 1C neuron across trials, grouped the 1C neurons by their preferred frequency, and selected a randomly chosen subset of 10 neurons from the same frequency group to drive each cortical neuron. (d) The top left plot shows the sound waveform presented in the 1C recordings used as input to the model cortical network. The top right plot shows PSTHs formed by averaging 1C responses across trials and across all 1C neurons in each preferred frequency group. The raster plots show the recorded responses of two cortical populations on successive trials, along with the activity generated by the network model fit to each recording when driven by 1C responses to the same sounds. (e)A scatter plot showing the noise correlations of responses measured from the actual recordings and from simulations of the network model fit to each recording when driven by 1C responses to the same sounds. The Spearman’s rank correlation for the recordings versus the model were p<10^−5^.

### Strong inhibition suppresses noise correlations

Our main interest was in understanding how the different network-level mechanisms that are capable of controlling intrinsic dynamics contribute to the correlated variability in responses evoked by sensory stimuli. The wide variety of intrinsic dynamics in our recordings was reflected in the differences in evoked responses across recording types; while some recordings contained strong, reliable responses to the onset of a stimulus, other recordings contained responses that were highly variable across trials (Figure 4a). There were also large differences in the extent to which the variability in evoked responses was correlated across the neurons in each recording; pairwise noise correlations were large in some recordings and extremely weak in others, even when firing rates were similar (Figure 4b).

Because evoked spike patterns can depend strongly on the specifics of the sensory stimulus, we could not make direct comparisons between experimental responses across different species and modalities; our goal was to identify the internal mechanisms that are responsible for the differences in noise correlations across recordings and, thus, any differences in spike patterns due to differences in external input would confound our analysis. To overcome this confound and enable the comparison of noise correlations across recording types, we simulated the response of the network to the same external input for all recordings. We constructed the external input using recordings of spiking activity from the inferior colliculus (IC), a primary relay nucleus in the sub-cortical auditory pathway (Figure 4c–d). Using the subset of our cortical recordings in which we presented the same sounds that were also presented during the IC recordings, we verified that the noise correlations in the simulated cortical responses were similar to those in the recordings (Figures 4e).

The parameter sweeps described in Figure 2 demonstrated that there are multiple features of the model network that can control its intrinsic dynamics, and a similar analysis of the noise correlations in simulated responses to external input produced similar results (Supplementary Figure 4).To gain insight into which of these features could account for the differences in noise correlations across our recordings, we examined the dependence of the strength of the noise correlations in each recording on each of the model parameters. While several parameters were able to explain a significant amount of the variance in noise correlations across recordings, the amount of variance explained by the strength of inhibitory feedback was by far the largest (Figure 5a). The predominance of inhibition in the control of noise correlations was confirmed by the measurement of partial correlations (the correlation between the noise correlations and each parameter that remains after factoring out the influence of the other parameters; partial *r*^2^ for inhibition: 0.67, excitation: 0.02, adaptation: 0.08, tonic input spread: 0.17, and tonic input baseline: 0.04). We also performed parameter sweeps to confirm that varying only the strength of inhibition was sufficient to result in large changes in noise correlations in the parameter regime of each recording (Figure 5b).

**Figure 5.**
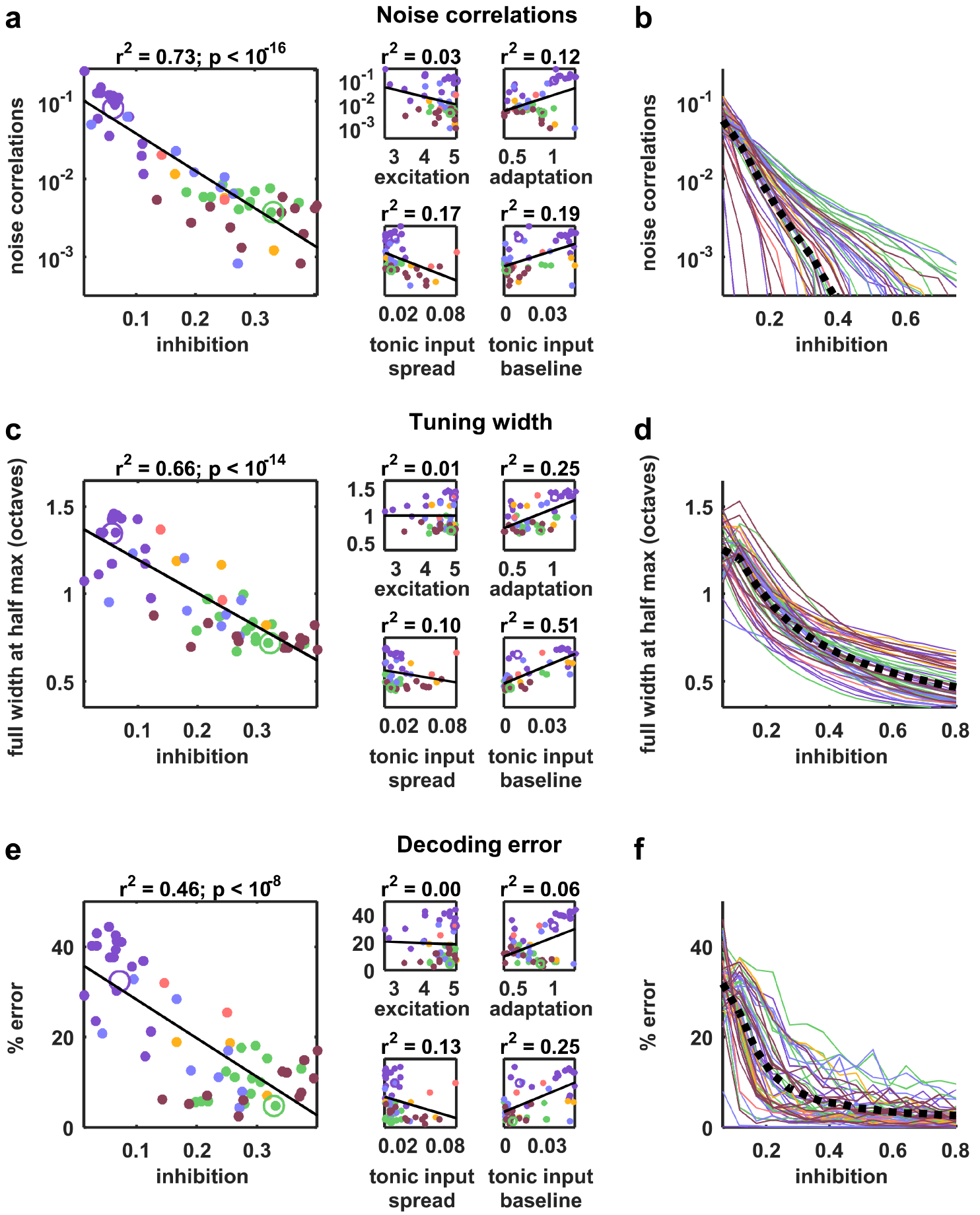
**Strong inhibition suppresses noise correlations and enhances selectivity and decoding** (a) Scatter plots showing the mean pairwise noise correlations measured from simulations of the network model fit to each recording when driven by external input versus the value of the different model parameters. Colors correspond to recording types as in Figure 4. The recordings shown in Figure 4d are denoted by open circles. Spearman’s rank correlation p-values for inhibition, excitation, adaptation, tonic input spread, and tonic input baseline were p<10^−18^, p=0.339, p=0.011, p<10^−2^, and p<10^−3^ respectively. (b) The mean pairwise noise correlations measured from network simulations with different values of the inhibition parameter w,. The values of all other parameters were held fixed at those fit to each recording. Each line corresponds to one recording. Colors correspond to recording types as in Figure 4. (c,e) Scatter plots showing tuning width and decoding error, plotted as in panel a. For panel c, Spearman’s rank correlation p-values for inhibition, excitation, adaptation, tonic input spread, and tonic input baseline were p<10^−15^, p=0.642, p<10^−4^, p<10^−2^, and p<10^−9^ respectively. For panel e, Spearman’s rank correlation p-values for inhibition, excitation, adaptation, tonic input spread, and tonic input baseline were p<10^−9^, p=0.799, p=0.0766, p<10^−2^, and p<10^−4^ respectively. (d,f) The tuning width and decoding error measured from network simulations with different values of the inhibition parameter w_I_ plotted as in panel b.

### Strong inhibition sharpens tuning and enables accurate decoding

We also examined how different features of the network controlled other aspects of evoked responses. We began by examining the extent to which differences in the value of each model parameter could explain differences in stimulus selectivity across recordings. To estimate selectivity, we drove the model network that was fit to each cortical recording with external inputs constructed from IC responses to tones, and used the simulated responses to measure the width of the frequency tuning curves of each model neuron. Although each model network received the same external inputs, the selectivity of the neurons in the different networks varied widely. The average tuning width of the neurons in each network varied most strongly with the strength of the inhibitory feedback in the network (Figure 5c; partial *r*^2^ for inhibition: 0.74, excitation: 0.06, adaptation: 0.48, tonic input spread: 0.01, and tonic input baseline: 0.37), and varying the strength of inhibition alone was sufficient to drive large changes in tuning width (Figure 5d). These results are consistent with experiments demonstrating that inhibition can control the selectivity of cortical neurons^49^, but suggest that this control does not require structured lateral inhibition.

We also investigated the degree to which the activity patterns generated by the model fit to each cortical recording could be used to discriminate different external inputs. We trained a decoder to infer which of seven possible stimuli evoked a given single-trial activity pattern and examined the extent to which differences in the value of each model parameter could account for the differences in decoder performance across recordings. Again, the amount of variance explained by the strength of inhibitory feedback was by far the largest (Figure 5e; partial *r*^2^ for inhibition: 0.5, excitation: 0.16, adaptation 0.27, tonic input spread 0.02, and tonic input baseline 0.03); decoding was most accurate for activity patterns generated by networks with strong inhibition, consistent with the weak noise correlations and high selectivity of these networks. Parameter sweeps confirmed that varying only the strength of inhibition was sufficient to result in large changes in decoder performance (Figure 5f).

### Activity of fast-spiking (FS) neurons is increased during periods of cortical desynchronization with weak noise correlations

Our model-based analyses suggest an important role for feedback inhibition in controlling the way in which responses to sensory inputs are shaped by intrinsic dynamics. In particular, our results predict that inhibition should be strong in dynamical regimes with weak noise correlations. To test this prediction, we performed further analysis of our recordings to estimate the strength of inhibition in each recorded population. We classified the neurons in each recording based on the width of their spike waveforms (Supplementary Figure 7). The waveforms for all recording types fell into two distinct clusters, allowing us to separate fast-spiking (FS) neurons from regular-spiking (RS) neurons. In general, more than 90% of FS cortical neurons have been reported to be parvalbumin-positive (PV+) inhibitory neurons^50–55^, and this value approaches 100% in the deep cortical layers where we recorded^56^. While the separation of putative inhibitory and excitatory neurons based on spike waveforms is imperfect (nearly all FS neurons are inhibitory, but a small fraction (less than 20%) of RS neurons are also inhibitory^57^), it is still effective for approximating the overall levels of inhibitory and excitatory activity in a population.

Given the results of our model-based analyses, we hypothesized that the overall level of activity of FS neurons should vary inversely with the strength of noise correlations. To identify sets of trials in each recording that were likely to have either strong or weak noise correlations, we measured the level of cortical synchronization. Previous studies have shown that noise correlations are strong when the cortex is in a synchronized state, where activity is dominated by concerted, large-scale fluctuations, and weak when the cortex is in a desynchronized state, where these fluctuations are suppressed^7,11^.

We began by analyzing our recordings from V1 of awake mice. We classified the cortical state during each stimulus presentation based on the ratio of low-frequency LFP power to high-frequency LFP power^58^ and compared evoked responses across the most synchronized and desynchronized subsets of trials (Figure 6a). As expected, noise correlations were generally stronger during synchronized trials than during desynchronized trials, and this variation in noise correlations with cortical synchrony was evident both within individual recordings and across animals (Figure 6b–c). As predicted by our model-based analyses, the change in noise correlations with cortical synchrony was accompanied by a change in FS activity; there was a four-fold increase in the mean spike rate of FS neurons from the most synchronized trials to the most desynchronized trials, while RS activity remained constant (Figure 6d–f).

We next examined our recordings from gerbil A1 under urethane in which the cortex exhibited transitions between distinct, sustained synchronized and desynchronized states (Figure 6g). As in our awake recordings, cortical desynchronization under urethane was accompanied by a decrease in noise correlations and an increase in FS activity (Figures 6h–k). In fact, both FS and RS activity increased with cortical desynchronization under urethane, but the increase in FS activity was much larger. The increase in RS activity suggests that cortical desynchronization under urethane may involve other mechanisms in addition to an increase in feedback inhibition (a comparison of the model parameters fit to desynchronized and synchronized urethane recordings (Supplementary Figure 3) suggests that the average level of tonic input is significantly higher during desynchronization (desynchronized: 0.075+/−0.008, synchronized: 0.0195+/−0.0054, *p* =0.006)).

Finally, we examined our remaining recordings from gerbil A1 under either ketamine/xylazine (KX) or fentanyl/medetomidine/midazolam (FMM) anesthesia. In these recordings, the cortex did not transition between different dynamical regimes, so we could not track changes in noise correlations and FS activity within individual recordings. However, recordings under KX and FMM exhibited stable states with strong and weak noise correlations respectively^7^ (Figure 7a), so we were able to make comparisons across recordings. Noise correlations under FMM were extremely weak, while those under KX were the largest in any of our recordings, so we expected FS activity under FMM to be much higher than that under KX. Surprisingly, our initial analysis suggested the opposite; the average spike rate of FS neurons under KX was larger than that under FMM (Figure 7b). Further analysis revealed, however, that there were many fewer FS neurons in our KX recordings than in our FMM recordings (Figure 7c), and this effect was most pronounced for FS neurons with low spike rates (Figure 7d). The low number of FS neurons in our KX recordings suggests that many FS neurons become completely silent under KX (all recordings were made in the same region of gerbil A1 with the same multi-tetrode arrays, so a similar number of FS neurons should be expected). When we measured FS activity as the sum of all spiking in each recording rather than the average spike rate of each neuron, the amount of FS activity was indeed much larger under FMM than under KX, consistent with our observations in other recording types and the predictions of our model-based analyses that stronger overall inhibition is accompanied by lower noise correlations (Figure 7e–f).

**Figure 6.**
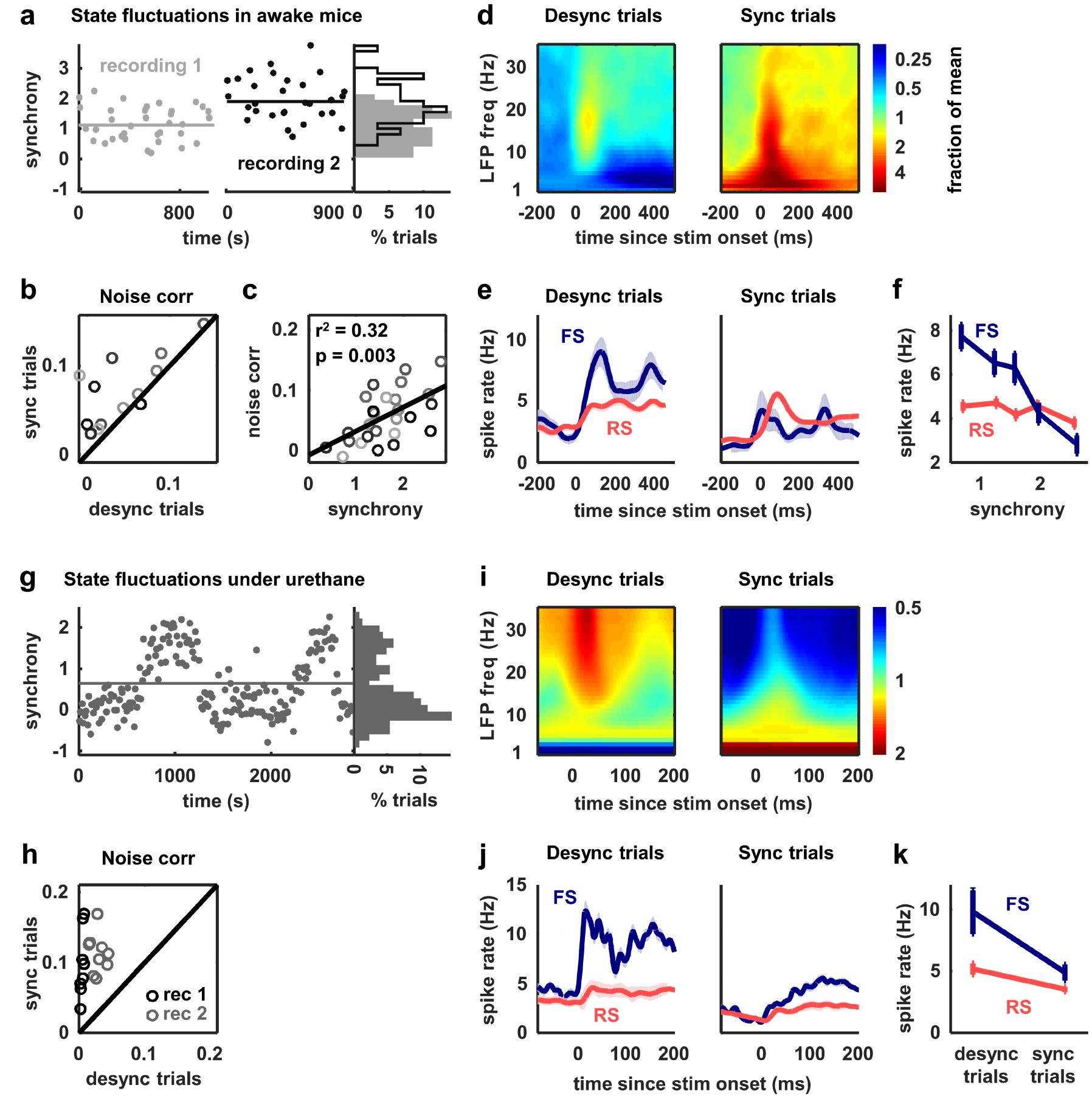
**Fast-spiking neurons are more active during periods of cortical desynchronization with weak noise correlations** (a) The cortical synchrony at different points during two recordings from VI of awake mice, measured as the log of the ratio of low-frequency (3 −10 Hz) LFP power to high-frequency (11 - 96 Hz). The distribution of synchrony values across each recording is also shown. The lines indicate the median of each distribution. (b) A scatter plot showing the noise correlations measured during trials in which the cortex was in either a relatively synchronized (‘sync’) or desynchronized (‘desync’) state for each recording. Each point indicates the mean pairwise correlations between the spiking activity of all pairs of neurons in one recording (after binning the activity in 15 ms bins). Trials with the highest 50% of synchrony values were classified as sync and trials with the lowest 50% of synchrony values were classified as desync. Values for 13 different recordings are shown. The Wilcoxon two-sided signed-rank test p-value was p<10^−2^. (c) A scatter plot showing noise correlations versus the mean synchrony for trials with the highest and lowest 50% of synchrony values for each recording. Colors indicate different recordings. The Spearman’s rank correlation significance among all recordings was p<10^−2^. (d) Spectrograms showing the average LFP power during trials with the highest (sync) and lowest (desync) 20% of synchrony values across all recordings. The values shown are the deviation from the average spectrogram computed over all trials. (e) The average PSTHs of FS and RS neurons measured from evoked responses during trials with the highest (sync) and lowest (desync) 20% of synchrony values across all recordings. The lines show the mean across all neurons, and the error bars indicate +/−1 SEM. (f) The average spike rate of FS and RS neurons during the period from 0 to 500 ms following stimulus onset, averaged across trials in each synchrony quintile. The lines show the mean across all neurons, and the error bars indicate +/−1 SEM. The Wilcoxon two-sided signed-rank test comparing FS activity between the highest and lowest quintile had a significance of p<10^−9^ and for RS activity, the significance was p<10^−2^. (g) The cortical synchrony at different points during a urethane recording, plotted as in panel a. The line indicates the value used to classify trials as synchronized (sync) or desynchronized (desync). (h)A scatter plot showing the noise correlations measured during trials in which the cortex was in either a synchronized (sync) or desynchronized (desync) state. Values for two different recordings are shown. Each point for each recording shows the noise correlations measured from responses to a different sound. The Wilcoxon two-sided signed-rank test between sync and desync state noise correlations had a significance of p<10 ^−3^. (i) Spectrograms showing the average LFP power during synchronized and desynchronized trials, plotted as in panel (j) The average PSTHs of FS and RS neurons during synchronized and desynchronized trials, plotted as in panel e. (k) The average spike rate of FS and regular-spiking RS neurons during the period from 0 to 500 ms following stimulus onset during synchronized and desynchronized trials. The points show the mean across all neurons, and the error bars indicate +/−1 SEM. The Wilcoxon two-sided signed-rank test comparing FS activity between the sync and desync had a significance of p<10^−3^ and for RS activity, the significance was p<10^−5^.

**Figure 7.**
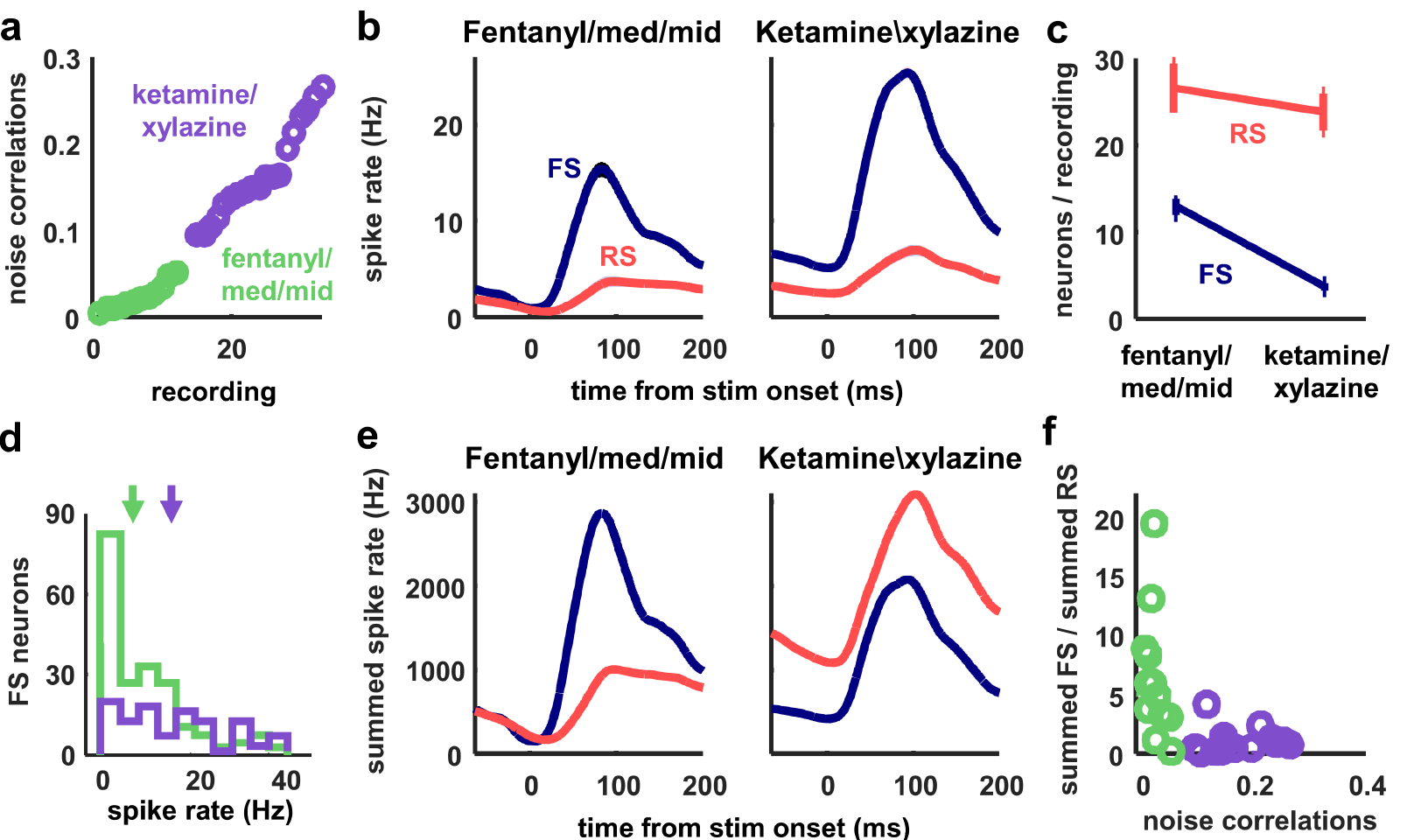
**Many fast-spiking neurons are silent under ketamine/xylazine anesthesia** (a) The noise correlations measured from recordings of responses to speech in gerbil A1 under ketamine/xylazine (KX) and fentanyl/medetomidine/midazolam (FMM). Each point indicates the mean pairwise correlations between the spiking activity of all pairs of neurons in one recording (after binning the activity in 15 ms bins). Recordings were sorted in order of increasing noise correlations. (b) The average PSTHs of FS and RS neurons under FMM or KX, plotted as in Figure 6. (c) The average number of FS and RS neurons in recordings under FMM and KX. The points show the mean across all recordings, and the error bars indicate +/−1SEM. The Wilcoxon two-sided rank-sum test between the number of FS neurons in FMM versus KX yielded a significance of p<10^−3^, and between the number of RS neurons, p=0.381. (d) Histograms of the average spike rate of FS neurons during the period from 0 to 500 ms following stimulus onset in recordings under FMM or KX. The arrows indicate the median values. (e) The summed PSTHs of FS and RS neurons under FMM or KX, plotted as in Figure 6. (f) The ratio of the total number of spikes from FS and RS neurons during the period from 0 to 500 msfollowing stimulus onset. Each point shows the value for one recording. The Spearman’s rank correlation was *r*^2^=0.31 and its significance was p<10^−2^.

## Discussion

We have shown here that a deterministic spiking network model is capable of intrinsically generating population-wide fluctuations in neural activity, without requiring external modulating inputs. Such fluctuations have been recently shown to arise in localized cortical networks, in both awake and anesthetized animals, and are not due to external inputs^9^. However, no models have been previously shown to realistically reproduce such coordinated activity in a local network of connected neurons. By fitting our spiking network model with adaptation currents directly to experimental recordings, we demonstrated that the model is able to reproduce the wide variety of multineuron cortical activity patterns observed in vivo without the need for external noise. Through chaotic amplification of small perturbations, the model generates activity with both trial-to-trial variability in the spike times of individual neurons and coordinated, large-scale fluctuations of the entire network. These fluctuations continue in the presence of sensory stimulation, thus creating noise correlations in a deterministic neural network.

The development of a network model that can reproduce experimentally-observed activity patterns through intrinsic variability alone is a major advance beyond previous models^20,24,29,35^. Networks in the classical balanced state produce activity with zero mean pairwise correlations between neurons^24, 33, 35^ and, thus, are not suitable to describe the population-wide fluctuations that are observed in many brain states in vivo^8^. To obtain single neuron rate fluctuations in balanced networks, structured connectivity has been used to create clustered networks^24^. However, while clustered networks do produce activity with positive correlations between a small fraction of neuron pairs (less than 1 in 1000), the average noise correlations across all pairs are still near zero and, thus, these networks are still unable to generate population-wide fluctuations.

We were able to overcome the limitations of previous models and generate intrinsic large-scale variability that is quantitatively similar to that observed in vivo by using spike-frequency adaptation currents in excitatory neurons, which have been well-documented experimentally^50,59^. The population-wide fluctuations generated by the interaction between recurrent excitation and adaptation were a robust feature of the network and persisted in more sophisticated networks that included multiple conductance timescales, many more neurons, spiking inhibitory neurons, structured connectivity, and kurtotic distributions of synaptic efficacies (see Supplementary Figure 2).

Although several features of the model network are capable of controlling its intrinsic dynamics, our analysis suggests that differences in feedback inhibition account for the differences in correlations across our in vivo recordings. When we fit the model to each of our individual recordings, we found that noise correlations, as well as stimulus selectivity and decoding accuracy, varied strongly with the strength of inhibition in the network. We also found that the activity of putative inhibitory neurons in our recordings was increased during periods of cortical desynchronization with weak noise correlations. Taken together, these results suggest that the control of correlated variability by inhibition plays a critical role in modulating the impact of intrinsic cortical dynamics on sensory responses.

### Inhibition controls the strength of the large-scale fluctuations that drive noise correlations

Our results are consistent with experiments showing that one global dimension of variability largely explains both the pairwise correlations between neurons^8^ and the time course of population activity^29^. In our network model, the coordinated, large-scale fluctuations that underlie this global dimension of variability are generated primarily by the interaction between recurrent excitation and adaptation. When inhibition is weak, small deviations from the mean spike rate can be amplified by strong, non-specific, recurrent excitation into population-wide events (up states). These events produce strong adaptation currents in each activated neuron, which, in turn, result in periods of reduced spiking (down states)^28,44,45,60^. The alternations between up states and down states have an intrinsic periodicity given by the timescale of the adaptation currents, but the chaotic nature of the network adds an apparent randomness to the timing of individual events, thus creating intrinsic temporal variability.

The intrinsic temporal variability in the network imposes a history dependence on evoked responses; because of the build-up of adaptation currents during each spiking event, external inputs arriving shortly after an up state will generally result in many fewer spikes than those arriving during a down state^28^. This history dependence creates a trial-to-trial variability in the total number of stimulus-evoked spikes that is propagated and reinforced across consecutive stimulus presentations to create noise correlations. However, when the strength of the inhibition in the network is increased, the inhibitory feedback is able to suppress some of the amplification by the recurrent excitation, and the transitions between clear up and down states are replaced by weaker fluctuations of spike rate that vary more smoothly over time. If the strength of the inhibition is increased even further, such that it becomes sufficient to counteract the effects of the recurrent excitation entirely, then the large-scale fluctuations in the network disappear, weakening the history dependence of evoked responses and eliminating noise correlations.

### Strong inhibition sharpens tuning curves and enables accurate decoding by stabilizing network dynamics

Numerous experiments have demonstrated that inhibition can shape the tuning curves of cortical neurons, with stronger inhibition generally resulting in sharper tuning^61^. The mechanisms involved are still a subject of debate, but this sharpening is often thought to result from structured connectivity that produces differences in the tuning of the excitatory and inhibitory synaptic inputs to individual neurons; lateral inhibition, for example, can sharpen tuning when neurons with similar, but not identical, tuning properties inhibit each other. Our results, however, demonstrate that strong inhibition can sharpen tuning in a network without any structured connectivity simply by controlling its dynamics.

In our model, broad tuning curves result from the over-excitability of the network. When inhibition is weak, every external input will eventually excite every neuron in the network because those neurons that receive the input directly will relay indirect excitation to the rest of the network. When inhibition is strong, however, the indirect excitation is largely suppressed, allowing each neuron to respond selectively to only those external inputs that it receives either directly or from one of the few other neurons to which it is strongly coupled. Thus, when inhibition is weak and the network is unstable, different external inputs will trigger similar population-wide events^62^, so the selectivity of the network in this regime is weak and its ability to encode differences between sensory stimuli is poor. In contrast, when inhibition is strong and the network is stable, different external inputs will reliably drive different subsets of neurons, and the activity patterns in the network will encode different stimuli with high selectivity and enable accurate decoding.

### Two different dynamical regimes with weak noise correlations

A number of studies have observed that the noise correlations in cortical networks can be extremely weak under certain conditions^5,7,35,63^. It was originally suggested that noise correlations were weak because the network was in an asynchronous state in which neurons are continuously depolarized with a resting potential close to the spiking threshold^33 35^. Experimental support for this classical asynchronous state has been provided by intracellular recordings showing that the membrane potential of cortical neurons is increased during locomotion^19^ and hyper-arousal^12^, resulting in tonic spiking. However, other experiments have shown that the membrane potential of cortical neurons in behaving animals can also be strongly hyperpolarized with clear fluctuations between up and down states^19,64–66^.

These apparently conflicting results suggest that there may be multiple dynamical regimes in behaving animals that are capable of producing weak noise correlations. There is mounting evidence suggesting that different forms of arousal may have distinct effects on neural activity^67^. While most forms of arousal tend to reduce the power of low-frequency fluctuations in membrane potential^19,66,68^, locomotion tends to cause a persistent depolarization of cortical neurons and drive tonic spiking. In contrast, task-engagement in stationary animals is generally associated with hyperpolarization and suppression of activity^15,18,19,66,69^. The existence of two different dynamical regimes with weak noise correlations was also apparent in our recordings; while some recordings with weak noise correlations resembled the classical asynchronous state with spontaneous activity consisting of strong, tonic spiking (e.g. desynchronized urethane recordings and some awake recordings), other recordings with weak noise correlations exhibited a suppressed state with relatively low spontaneous activity that contained clear, albeit weak, up and down states (e.g. FMM recordings and other awake recordings). Our model was able to accurately reproduce spontaneous activity patterns and generate evoked responses with weak noise correlations in both of these distinct regimes.

In addition to strong inhibition, the classical asynchronous state with strong, tonic spiking appears to require a combination of weak adaptation and an increase in the number of neurons receiving strong tonic input (see parameter sweeps in Figures 2c–d and parameter values for awake mouse V1 recordings in Supplementary Figure 4). Since large-scale fluctuations arise from the synchronization of adaptation currents across the population, reducing the strength of adaptation diminishes the fluctuations^28,45,60^. Increasing tonic input also diminishes large-scale fluctuations, but in a different way^44^; when a subset of neurons receive increased tonic input, their adaptation currents may no longer be sufficient to silence them for prolonged periods, and the activity of these neurons during what would otherwise be a down state prevents the entire population from synchronizing. When the network in the asynchronous state is driven by an external input, it responds reliably and selectively to different inputs. Because the fluctuations in the network are suppressed and its overall level of activity remains relatively constant, every input arrives with the network in the same moderately-adapted state, so there is no history dependence to create noise correlations in evoked responses.

Unlike in the classical asynchronous state, networks in the suppressed state have slow fluctuations in their spontaneous activity, and the lack of noise correlations in their evoked responses is due to different mechanisms (see parameter values for gerbil A1 FMM recordings in Supplementary Figure 4). The fluctuations in the hyperpolarized network are only suppressed when the network is driven by external input. In our model, this suppression of the correlated variability in evoked responses is caused by the supralinearity of the feedback inhibition^39^. The level of spontaneous activity driven by the tonic input to each neuron results in feedback inhibition with a relatively low gain, which is insufficient to suppress the fluctuations created by the interaction between recurrent excitation and adaptation. However, when the network is strongly driven by external input, the increased activity results in feedback inhibition with a much higher gain, which stabilizes the network and allows it to respond reliably and selectively to different inputs. This increase in the inhibitory gain of the driven network provides a possible mechanistic explanation for the recent observation that the onset of a stimulus quenches variability^70^ and switches the cortex from a synchronized to a desynchronized state^65^, as well as the suppression of responses to high-contrast stimuli in alert animals^71^.

### Experimental evidence for inhibitory stabilization of cortical dynamics

The results of several previous experimental studies also support the idea that strong inhibition can stabilize cortical networks and enhance sensory coding. In vitro studies have shown that pharmacologically reducing inhibition increases the strength of the correlations between excitatory neurons in a graded manner^72^. In vivo whole-cell recordings in awake animals have demonstrated that the stimulus-evoked inhibitory conductance is much larger than the corresponding excitatory conductance^43^. This strong inhibition in awake animals quickly shunts the excitatory drive and results in sharper tuning and sparser firing than the balanced excitatory and inhibitory conductances observed under anesthesia. While some of the increased inhibition in awake animals may be due to inputs from other brain areas^73^, the increased activity of local inhibitory interneurons appears to play an important role^16,74, 75^. However, not all studies have observed increased inhibition in behaving animals^76^, and the effects of behavioral state on different inhibitory interneuron types are still being investigated^66, 77, 78^.

The effects of local inhibition on sensory coding have also been tested directly using optogenetics. While the exact roles played by different inhibitory neuron types are still under investigation^79,80^, the activation of inhibitory interneurons generally results in sharper tuning, weaker correlations, and enhanced behavioral performance^49,81,82^, while suppression of inhibitory interneurons has the opposite effect, decreasing the signal-to-noise ratio and reliability of evoked responses across trials^82,83^. These results demonstrate that increased inhibition enhances sensory processing and are consistent with the overall suppression of cortical activity that is often observed during active behaviors^15,16, 69, 75^. In fact, one recent study found that the best performance in a detection task was observed on trials in which the pre-stimulus membrane voltage was hyperpolarized and low-frequency fluctuations were absent^19^, consistent with a suppressed, inhibition-stabilized network state.

### Neuromodulators and inhibitory control of cortical dynamics

Neuromodulators can exert a strong influence on cortical dynamics by regulating the balance of excitation and inhibition in the network. While the exact mechanisms by which neuromodulators control cortical dynamics are not clear, several lines of evidence suggest that neuromodulator release serves to enhance sensory processing by increasing inhibition. Increases in acetylcholine (ACh) and norepinephrine (NE) have been observed during wakefulness and arousal^84,85^, and during periods of cortical desynchronization in which slow fluctuations in the LFP are suppressed^82,86,87^. Stimulation of the basal forebrain has been shown to produce ACh-mediated increases in the activity of FS neurons and decrease the variability of evoked responses in cortex^86–88^. In addition, optogenetic activation of cholinergic projections to cortex resulted in increased firing of SOM+ inhibitory neurons and reduced slow fluctuations^82^. The release of NE in cortex through microdialysis had similar effects, increasing fast-spiking activity and reducing spontaneous spike rates^87^, while blocking NE receptors strengthened slow fluctuations in membrane potential^12^. More studies are needed to tease apart the effects of different neurotransmitters on pyramidal neurons and interneurons^82,87,88^, but most of the existing evidence is consistent with our results in suggesting that neuromodulators can suppress intrinsic fluctuations and enhance sensory processing in cortical networks by increasing inhibition.

### Simulating the neocortical architecture

Recently, there have been major efforts toward constructing neural network simulations of increasingly larger scale^89^ and biological fidelity^90^. There are many biological sources of information that can constrain the parameters of such large-scale simulations, including physiological^90^, anatomical^91–93^ and genetic^94,95^. However, while such complex simulations may be able to capture the relevant properties of a circuit and replicate features of its neural activity in detail, they may not necessarily provide direct insight into the general mechanisms that underlie the circuit’s function. Thus, a complementary stream of research is needed to seek minimal functional, yet physiologically-based, models that are capable of reproducing relevant phenomena. The model we have investigated here includes only a very restricted set of physiological properties, yet is able to reproduce the full range of dynamics observed across different species, brain areas, and behavioral states. This simple model provides a compact and intuitive description of the circuit mechanisms that are capable of coordinated dynamics in networks with intrinsic variability. We have already shown that the same mechanisms can also control the dynamics of more complex functional models, but further work is needed to develop methods to bridge the gap between functional models and large-scale digital reconstructions.

## Methods

All of the recordings analyzed in this study have been described previously. Only a brief summary of the relevant experimental details are provided here. Each recording is considered as a single sample point to which we fit our model. Thus, our sample size is 59. This is justified as sufficient because our samples span multiple brain regions and multiple species, and may be considered as representative activity for a range of different brain states. Due to the sample size, we used the Spearman’s (non-parametric) rank correlation in most of our analyses.

### Mouse V1

The experimental details for the mouse V1 recordings have been previously described^8^. The recordings were performed on male and female mice older than 6–7 weeks, of C57BL/6J strain. Mice were on 12 hours non-reversed light-dark regime. The mice were implanted with head plates under anaesthesia. After head-plate implant each mouse was housed individually. After a few days of recovery the mice were accustomed to having their head fixed while sitting or standing in a custom built tube. On the day of the recording, the mice were briefly anaesthetised with isoflurane, and a small craniectomy above V1 was made. Recordings were performed at least 1.5h after the animals recovered from the anaesthesia. Buzsaki32 or A4x8 silicon probes were used to record the spiking activity of populations of neurons in the infragranular layers of V1.

Visual stimuli were presented on two of the three available LCD monitors, positioned 25 cm from the animal and covering a field of view of 120° × 60°, extending in front and to the right of the animal. Visual stimuli consisted of multiple presentations of natural movie video clips. For recordings of spontaneous activity, the monitors showed a uniform grey background.

### Rat A1

The experimental procedures for the rat A1 recordings have been previously described^96^. Briefly, head posts were implanted on the skull of male Sprague Dawley rats (300–500 g, normal light cycle, regular housing conditions) under ketamine-xylazine anesthesia, and a hole was drilled above the auditory cortex and covered with wax and dental acrylic. After recovery, each animal was trained for 6–8 d to remain motionless in the restraining apparatus for increasing periods (target, 1–2 h). On the day of the recording, each animal was briefly anesthetized with isoflurane and the dura resected; after a 1 h recovery period, recording began. The recordings were made from infragranular layers of auditory cortex with 32-channel silicon multi-tetrode arrays.

Sounds were delivered through a free-field speaker. As stimuli we used pure tones (3, 7, 12, 20, or 30 kHz at 60 dB). Each stimulus had duration of 1s followed by 1s of silence. All procedures conformed to the National Institutes of Health Guide for the Care and Use of Laboratory Animals.

### Gerbil A1

The gerbil A1 recordings have been described in detail previously^7^. Briefly, adult male gerbils (70–90 g, P60–120, normal light-dark cycle, group housed) were anesthetized with one of three different anesthetics: ketamine/xylazine (KX), fentanyl/medetomidine/midazolam (FMM), or urethane. A small metal rod was mounted on the skull and used to secure the head of the animal in a stereotaxic device in a sound-attenuated chamber. A craniotomy was made over the primary auditory cortex, an incision was made in the dura mater, and a 32-channel silicon multi-tetrode array was inserted into the brain. Only recordings from A1 were analyzed. Recordings were made between 1 and 1.5 mm from the cortical surface (most likely in layer V). All gerbils were used in this study, except for one gerbil under FMM which exhibited little to no firing over the recordings.

Sounds were delivered to speakers coupled to tubes inserted into both ear canals for diotic sound presentation along with microphones for calibration. Repeated presentations of a 2.5 s segment of human speech were presented at a peak intensity of 75 dB SPL. For analyses of responses to different speech tokens, seven 0.25 s segments were extracted from the responses to each 2.5 s segment.

### Gerbil IC

The gerbil IC recordings have been described in detail previously^97^. Recordings were made under ketamine/xylazine anesthesia using a multi-tetrode array placed in the low-frequency laminae of the central nucleus of the IC. Experimental details were otherwise identical to those for gerbil A1. In addition to the human speech presented during the A1 recordings, tones with a duration of 75 ms and frequencies between 256 Hz and 8192 Hz were presented at intensities between 55 and 85 dB SPL with a 75 ms pause between each presentation.

All relevant data are available from the authors upon request.

### Spiking network model

We developed a network model using conductance-based quadratic integrate and fire neurons. There are three currents in the model: an excitatory, an inhibitory and an adaptation current. The subthreshold membrane potential for a single neuron *i* obeys the equation

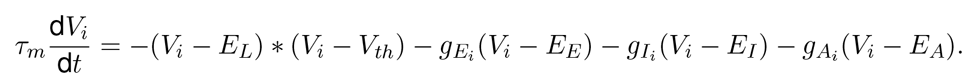

When *V* > *V_th_*, a spike is recorded in the neuron and the neuron’s voltage is reset to *V_reset_* = 0 .9*V_th_*. For simplicity, we set *V_th_* = 1 and the leak voltage *E_L_* = 0. The excitatory voltage *E_E_* = 2 *V_th_* and *E_I_* = *E_A_* = −0.5*V_th_*. Each of the conductances has a representative differential equation which is dependent on the spiking of the neurons in the network at the previous time step, S_*t*−1_.

The excitatory conductance obeys

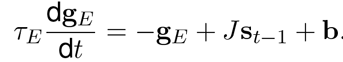

where *J* is the matrix of excitatory connectivity and b is the vector of tonic inputs to the neurons. The matrix of connectivity is random with a probability of 5% for the network of 512 neurons and their connectivities are randomly chosen from a uniform distribution between 0 and *w_E_*. The tonic inputs b have a minimum value *b*_0_, which we call the tonic input baseline added to a random draw from an exponential distribution with mean *b_1_*, which we call the tonic input spread, such that for neuron *i*, b(*i*) = *b*_0_ + exprnd(*b_l_*). The inhibitory conductance obeys

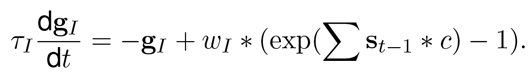

where *c* controls the gain of the inhibitory conductance.

The adaptation conductance obeys

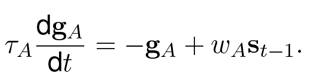

The simulations are numerically computed using Euler’s method with a time-step of 0.75 ms (this was the lock-out window used for spike-sorting the in vivo recordings and allowed for fast simulations). To avoid numerical instabilities at low voltages, we rectified the voltages at the activation potential of the inhibitory conductance. Each parameter set was simulated for 900 seconds. The timescales are set to *τ_m_* =20 ms, *τ_E_* =5.10 ms, *τ_I_* = 3.75 ms, *τ_A_* = 375 ms, and the inhibitory non-linearity controlled by *c* = 0.25. The remaining five parameters (*w_I_, w_A_, w_E_*, *b_l_*, and *b_0_*) were fit to the spontaneous activity from multi-neuron recordings using the techniques described below. Their ranges were (0.01–0.4), (0.4–1.45), (2.50–5.00), (0.005–0.10), and (0.0001–0.05) respectively.

To illustrate the ability of the network to generate activity patterns with macroscopic variability, we simulated spontaneous activity with a parameter set that produces up and down state dynamics. Figure 2a shows the membrane potential of a single neuron in this simulation and its conductances at each time step. Figure 2b shows the model run twice with the same set of initial conditions and parameters, but with an additional single spike inserted into the network on the second run (circled in green).

This code will be made available for use after publication.

### Parameter sweep analysis

Figure 2c and d summarize the effects of changing each parameter on the structure of the spontaneous activity patterns generated by the model. We held the values for all but one parameter fixed and swept the other parameter across a wide range of values. The fixed parameter values were set to approximately the median values obtained from fits to all in vivo recordings. Figure 5b, d, and f and Supplementary Figure 6 show the results of similar parameter sweep analyses for stimulus-driven activity with the external input to the network derived from IC activity as described below. For these analyses, the values of the parameters that were not swept were fixed at those fit to each individual recording.

### GPU implementation

We accelerated the network simulations by programming them on graphics processing units (GPUs) such that we were able to run them at 650x real time with 15 networks running concurrently on the same GPU. We were thus able to simulate ≈10000 seconds of simulation time in 1 second of real time. To achieve this acceleration, we took advantage of the large memory bandwidth of the GPUs. For networks of 512 neurons, the state of the network (spikes, conductances and membrane potentials) can be stored in the very fast ‘shared memory’ available on each multiprocessor inside a GPU. A separate network was simulated on each of the 8 or 15 multiprocesssors available (video cards were GTX 690 or Titan Black). Low-level CUDA code was interfaced with Matlab via mex routines.

### Summary statistics

Several statistics of spikes were used to summarize the activity patterns observed in the in vivo recordings and in the network simulations. Because there were on the order of 50 neurons in each recording, all of the statistics below were influenced by small sample effects. To replicate this bias in the analysis of network simulations, we subsampled 50 neurons from the network randomly and computed the same statistics we computed from the in vivo recordings.

The noise correlations between each pair of neurons in each recording were measured from responses to speech. The response of each neuron to each trial was represented as a binary vector with 15 ms time bins. The total correlation for each pair of neurons was obtained by computing the correlation coefficient between the actual responses. The signal correlation was computed after shuffling the order of repeated trials for each time bin. The noise correlation was obtained by subtracting the signal correlation from the total correlation.

The multi-unit activity (MUA) was computed as the sum of spikes in all neurons in bins of 15 ms.

The autocorrelation function of the MUA at time-lag *τ* was computed from the formula

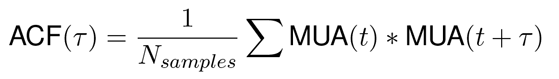

To measure the autocorrelation timescale, we fit one side of the ACF with a parametric function

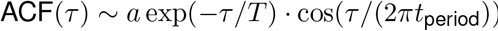

where *a* is an overall amplitude, *T* is a decay timescale and *t_period_* is the oscillation period of the autocorrelation function. There was not always a significant oscillatory component in the ACF, but the timescale of decay accurately captured the duration over which the MUA was significantly correlated.

### Parameter searches

To find the best fit parameters for each individual recording, we tried to find the set of model parameters for which the in vivo activity and the network simulations had the same statistics. We measured goodness of fit for each of the three statistics: pairwise correlations, the MUA distribution, the MUA ACF. Each statistic was normalized appropriately to order 1, and the three numbers obtained were averaged to obtain an overall goodness of fit.

The distance measure *D_c_* between the mean correlations *c_θ_* obtained from a set of parameters *θ* and the mean correlations *c_n_* in recording *n* was simply the squared error *D_c_(c_n_, c_θ_)* = (*c_n_ - c_θ_*)^2^. This was normalized by the variance of the mean correlations across recordings to obtain the normalized correlation cost Cost_*c*_, where 〈*x_n_*〉_*n*_ is used to denote the average of a variable *x* over recordings indexed by *n*.

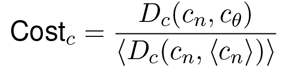

The distance measure *D_m_* for the MUA distribution was the squared difference summed over the order rank bins *k* of the distribution 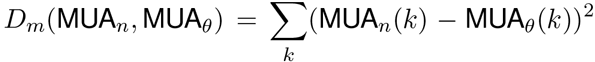. This was normalized by the distance between the data MUA and the mean data MUA. In other words, the cost measures how much closer the simulation is to the data distribution than the average of all data distributions.

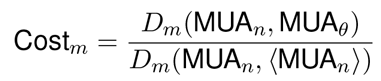

Finally, the distance measure *D_a_* for the autocorrelation function of the MUA was the squared difference summed over time lag bins *t* of the distribution 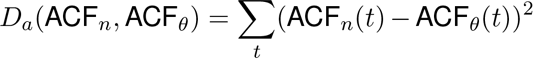.

This was normalized by the distance between the data ACF and the mean data ACF.

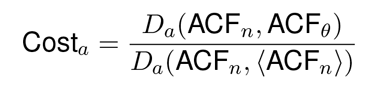

The total cost of parameters *θ* on recording *n* is therefore Cost(*n, θ*) = Cost_*c*_ + Cost_*m*_ + Cost_*a*_. Approximately one million networks were simulated on a grid of parameters for 600 seconds each of spontaneous activity, and their summary statistics (*c_θ_* MUA_*θ*_, and ACF_*θ*_) were retained. The Cost was smoothed for each recording by averaging with the nearest 10 other simulations on the grid. This ensured that some of the sampling noise was removed and parameters were estimated more robustly. The best fit set of parameters was chosen as the minimizer of this smoothed cost function, on a recording by recording basis.

### Alternative Gibbs sampling parameter optimization

We also demonstrate an alternative approach to finding the best fitting parameters through a sampling-based optimization procedure (Supplementary Figure 3). This reduces the necessary number of simulations from 1 million to 100,000. Future work might in principle devise even faster optimizations, thus allowing analysis on a bigger scale than presented here. Briefly, the sampling-based optimization is based on defining the energy landscape as the negative of the cost function, and thus defining a probability distribution over parameters P(*θ*) = *exp*(−Cost(*θ*)/*T*), where *T* is the “temperature”. We use a proposal distribution that always proposes neighbors of the current sample on the grid on which we did the full parameter sweeps, and accept the proposals according to the balance equations of Markov Chain Monte Carlo sampling (MCMC):

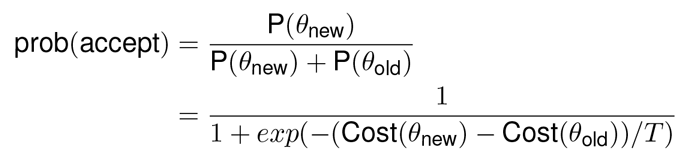

To avoid the MCMC chains getting stuck into low probability parts of the energy landscape, we restart the chain every 50 samples from the pool of already-sampled points, chosen with probability proportional to its P(*θ*). Furthermore, we allow the chain used to optimize the model parameters for one recording to use information from the chains used for the other recordings by pooling together the already-sampled points from all datasets and restarting chains based on all these points.

### NMDA and GABA_B_ conductance network

We added long timescale excitatory and inhibitory conductances to the model and simulated the model at multiple levels of inhibitory feedback strength. The strength of the NMDA conductance was 4% the strength of the AMPA conductance and *τ_NMDA_* = 100 ms (thus the integrated current was approximately the same as the AMPA integrated current injection). The strength of the GABA_b_ conductance was 2% the strength of the GABA conductance and it had the same timescale as NMDA. The parameter set used for Supplementary Figure 2a was *θ* = (0.51, *w_I_*, 2.6, 0.008, 0.037), where *w_I_* ranged from 0.02 to 0.25.

### Clustered neuronal network with intrinsic variability and spiking inhibitory neurons

We also simulated a clustered architecture with variability and adaptation currents. This model consisted of 144 clusters, each with 32 neurons, 8 of which were inhibitory neurons and 24 of which were excitatory neurons. The probability of within cluster excitatory-excitatory (E-E) connectivity was 0.3, and within cluster inhibitory-excitatory (I-E) and excitatory-inhibitory (E-I) were 0. 15 and 0.1 respectively. The probability of out of cluster E-E, I-E, and E-I connectivity were 0. 012, 0.03, and 0.01 respectively. The inhibitory-inhibitory connectivity was unclustered. The probability of connection was 0.01 and its strength was 0.17. The average connection strengths for E-E and I-E were 0.024 and 0.016 respectively. The E-I strength in Supplementary Figure 2b ranged from 0.025 to 0.057. The adaptation current had strength 0.45 and *τ_A_* = 220 ms. The membrane timescale for excitatory and inhibitory neurons were 25 ms and 5 ms respectively, and *τ_E_* = 6 ms and *τ_I_* = 3 ms.

### Stimulus-driven activity

Once the simulated networks were fit to the spontaneous neuronal activity, we drove them with an external input to study their evoked responses. The stimulus was either human speech (as presented during our gerbil A1 recordings) or pure tones. The external input to the network was constructed using recordings from 563 neurons from the inferior colliculus (IC). For all recordings in the IC the mean pairwise noise correlations were near-zero and the Fano Factors of individual neurons were close to 1^97^, suggesting that responses of IC neurons on a trial-by-trial basis are fully determined by the stimulus alone, up to Poisson-like variability. Thus, we averaged the responses of IC neurons over trials and drove the cortical network with this trial-averaged IC activity. We binned IC neurons by their preferred frequency in response to pure tones, and drove each model cortical neuron with a randomly chosen subset of 10 neurons from the same preferred-frequency bin. We rescaled the IC activity so that the input to the network had a mean value of 0.06 and a maximum value of 0.32, which was three times greater than the average tonic input.

We kept the model parameters fixed at the values fit to spontaneous activity and drove the network with 330 repeated presentations of the stimulus. We then calculated the statistics of the evoked activity. Noise correlations were measured in 15-ms bins as the residual correlations left after subtracting the mean response of each neuron to the stimulus across trials:

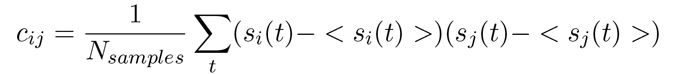

where *s_i_(t)* is the summed spikes of neuron i in a 15-ms bin and < *s_i_(t*) > is the mean response of neuron i to the stimulus. The noise correlation value given for each recording is the mean of *c_ij_*.

### Tuning width

To determine tuning width to sound frequency, we used responses of IC neurons to single tones as inputs to the model network. The connections from IC to the network were the same as described in the previous section. Because the connectivity was tonotopic and IC responses are strongly frequency tuned, the neurons in the model network inherited the frequency tuning. We did not model the degree of tonotopic fan-out of connections from IC to cortex and, as a result, the tuning curves of the model neurons were narrow relative to those observed in cortical recordings^7^. We chose the full width of the tuning curve at half-max as a standard measure of tuning width.

### Decoding tasks

We computed decoding error for a classification task in which the single-trial activity of all model neurons was used to infer which of seven different speech tokens was presented. The classifier was built on training data using a linear discriminant formulation in which the Gaussian noise term was replaced by Poisson likelihoods. Specifically, the activity of a neuron for each 15-ms bin during the response to each token was fit as a Poisson distribution with the empirically-observed mean. To decode the response to a test trial, the likelihood of each candidate token was computed and the token with the highest likelihood was assigned as the decoded class. This classifier was chosen because it is very fast and can be used to model Poisson-like variables, but we also verified that it produced decoding performance as good as or better than classical high-performance classifiers like support vector machines.

### Classifying FS and RS neurons

We classified fast-spiking and regular-spiking neurons based on their spike shape^8^. We determined the trough-to-peak time of the mean spike waveform after smoothing with a gaussian kernel of σ = 0.5 samples. The distribution of the trough-to-peak time *τ* was clearly bimodal in all types of recordings. Following^8^ we classified FS neurons in the awake data with *τ* < 0.6ms and RS neurons with *τ* > 0.8ms. The distributions of *τ* in the anesthetized data, although bimodal, did not have a clear separation point, so we conservatively required *τ* < 0.4ms to classify an FS cell in these recordings and *τ* > 0.65ms to classify RS neurons (see Supplementary Figure 7). The rest of the neurons were not considered for the plots in Figures 7 and 8 and are shown in gray on the histogram in Supplementary Figure 7.

Although one recent study has raised doubts on the accuracy of spike-width based classification^98^, a large number of other studies have shown 90–100% classification accuracy of FS neurons as PV-positive interneurons, and even^98^ show that the classification is near-perfect using other features of the spike waveform. We propose that the difference arises due to different properties of the bandpass filters used in these studies, and we show that in our datasets, trough-to-peak duration of the raw waveform is a highly bimodal distribution in all cases (see Supplementary Figure 7), unlike the distributions shown in^98^.

### Local field potential

The low-frequency potential (LFP) was computed by low-pass filtering the raw signal with a cutoff of 300 Hz. Spectrograms with adaptive time-frequency resolution were obtained by filtering the LFP with Hamming-windowed sine and cosine waves and the spectral power was estimated as the sum of their squared amplitudes. The length of the Hamming-window was designed to include two full periods of the sine and cosine function at the respective frequency, except for frequencies of 1 Hz and above 30 Hz, where the window length was clipped to a single period of the sine function at 1 Hz and two periods of the sine function at 30 Hz respectively. The synchrony level was measured as the log of the ratio of the low to high frequency power (respective bands: 3–10 Hz and 11–96 Hz, excluding 45–55 Hz to avoid the line noise). We did not observe significant gamma power peaks except for the line noise, in either the awake or anesthetized recordings.

### Dividing trials by synchrony

We computed a synchrony value for each trial in the 500-ms window following stimulus onset. For urethane recordings, the values had a clear bimodal distribution and we separated the top and bottom of these distributions into synchronized and desynchronized trials respectively. For awake recordings, the synchrony index was not clearly bimodal, but varied across a continuum of relatively synchronized and desynchronized states. To examine the effect of synchrony on noise correlations, we sorted all trials by their synchrony value, classified the 50% of trials with the lowest values as desynchronized and the 50% of trials with the highest values as synchronized, and computed the noise correlations for each set of trials for each recording. To examine the effect of synchrony on FS and RS activity, we pooled all trials from all recordings, divided them into quintiles by their synchrony value, and computed the average spike rates of FS and RS neurons for each set of trials. For Figures 6 and 7, noise correlations were computed aligned to the stimulus onsets in windows of 500ms, to match the window used for measuring FS and RS activity as well as LFP power.

